# *Zymoseptoria tritici* suppresses the host immune response and facilitates the success of avirulent strains in mixed infections

**DOI:** 10.1101/2023.05.13.540507

**Authors:** Alessio Bernasconi, Cécile Lorrain, Priska Flury, Julien Alassimone, Bruce A. McDonald, Andrea Sánchez-Vallet

## Abstract

Plants interact with a plethora of pathogenic microorganisms in nature. Pathogen-plant interaction experiments focus mainly on single-strain infections, typically ignoring the complexity of multi-strain infections even though mixed infections are common and critical for the infection outcome. The wheat pathogen *Zymoseptoria tritici* forms highly diverse fungal populations in which several pathogen strains often colonize the same leaf. Despite the importance of mixed infections, the mechanisms governing interactions between a mixture of pathogen strains within a plant host remain largely unexplored. Here we demonstrate that avirulent pathogen strains benefit from being in mixed infections with virulent strains. We show that virulent strains suppress the wheat immune response, allowing the avirulent strain to colonize the apoplast and to reproduce. Our experiments indicate that virulent strains in mixed infections can challenge the plant immune system both locally and systemically, providing a mechanistic explanation for the persistence of avirulent pathogen strains in fields planted to resistant host plants.

## Introduction

Complex biotic interactions within a plant can modulate the outcome of pathogen infections (Tollenaere *et al*., 2016, 2017). The dynamics of pathogen-host interactions involving mixed microbial infections can include cooperation, coexistence, and competition, where each infecting microbe may affect directly or indirectly the performance of the others (Abdullah *et al*., 2017). Co-infecting microbes can compete directly through the secretion of molecules (e.g., antimicrobial metabolites) that affect the fitness of the competitors (Rovenich *et al*., 2014; Tollenaere *et al*., 2016). Within a shared host, microorganisms can also interact indirectly when the performance of each interacting microbe depends on the response of the host (Alizon and Van Baalen, 2008; Mideo, 2009; Holt and Bonsall, 2017). For instance, plant susceptibility or resistance triggered by a pathogen can affect the success of coinfecting microbes. In fact, infections by the fungi *Fusarium oxysporum* and *Zymoseptoria tritici* predispose wheat to be colonized by the bacteria *Pseudomonas fluorescens* and *Pseudomonas syringae*, respectively, through the suppression of host antimicrobial substances (Notz *et al*., 2002; Seybold *et al*., 2020). The distinctive plant immune responses to pathogens with different lifestyles have been well documented across various systems (Glazebrook, 2005). Since plants are constantly challenged by an enormous diversity of pathogenic microorganisms with varying infection strategies (Gold *et al*., 2009; Laine, 2011; Susi *et al*., 2015; Barrett *et al*., 2021), it is reasonable to expect that genotype-specific host immune responses may affect the infection outcome of co-infecting pathogen strains.

Comprehensive studies of plant immune responses to pathogen infections have mostly been conducted using experimental systems in which a single host genotype is infected by a single pathogen strain. Virulent pathogen strains colonize the host by secreting effectors that suppress host immunity (He *et al*., 2004; Martin and Kamoun, 2012; Jiang *et al*., 2013; Gimenez-Ibanez *et al*., 2014; Lo Presti *et al*., 2015; Sánchez-Vallet, Mesters, *et al*., 2015; Fiorin *et al*., 2018; Haueisen *et al*., 2019, 2020). To counteract pathogen colonization, plants evolved the ability to specifically recognize certain forms of effectors - or avirulence factors (Avrs) – through the production of resistance (R) proteins, and subsequently trigger an immune response that hinders the progression of the pathogen (Jones and Dangl, 2006; Kanyuka and Rudd, 2019). Pathogen strains harboring recognized Avrs are known as avirulent strains (Flor, 1971; Bent and Mackey, 2008). The host immune response can act locally, occurring only where the avirulent strain interacts with the host, but it can also spread from the infection site to produce a systemic response (McDowell and Dangl, 2000; Vlot *et al*., 2021). In mixed infections, virulent and avirulent pathogen strains may co-exist and affect the outcome of infection. In particular, the induction of host immunity caused by avirulent pathogen strains might suppress the development of co-infecting virulent strains. It is also possible that suppression of the host immune response by virulent pathogen strains might facilitate infection by co-infecting avirulent strains (Barrett *et al*., 2021). Our experiments aimed to better understand the balance between these processes.

*Z. tritici* is a genetically diverse fungal wheat pathogen in which several distinct strains typically coinfect the same leaf on a plant (Linde *et al*., 2002; Zhan *et al*., 2005; Barrett *et al*., 2021; McDonald *et al*., 2022). *Z. tritici* hyphae penetrate through the stomata and grow in the apoplast without producing symptoms for a period that lasts between 10 and 14 days under controlled conditions. Subsequently, it produces necrotic lesions on wheat leaves and forms both asexual (pycnidia) and sexual (pseudothecia) reproductive structures (Kema *et al*., 1996; Duncan and Howard, 2000; Sánchez-Vallet, McDonald, *et al*., 2015). The septoria tritici blotch (STB) disease, caused by *Z. tritici,* leads to significant losses in wheat production worldwide and is managed mainly using host resistance and fungicides. Genetic control of STB is based on wheat resistance genes, of which 22 have been identified so far (Brading *et al*., 2002; Brown *et al*., 2015; Yang *et al*., 2018; Saintenac *et al*., 2021) and two (Stb6 and Stb16) have been cloned and functionally characterized (Saintenac *et al*., 2018, 2021). Stb6 recognizes avirulent isoforms of the fungal effector AvrStb6, following the gene-for-gene interaction model (Zhong *et al*., 2017; Kema *et al*., 2018), leading to the induction of an immune response that prevents the progression and asexual reproduction of AvrStb6-expressing strains (Zhong *et al*., 2017; Kema *et al*., 2018; Saintenac *et al*., 2018). Field populations of *Z. tritici* carry a diverse array of AvrStb6 isoforms, and virulent and avirulent strains typically coexist in the same field (Brunner and McDonald, 2018) and are capable of reproducing sexually (Kema *et al*., 2018). It has yet to be determined if and how mixed infections affect asexual reproduction and the persistence of avirulent strains.

In this work, we demonstrate that an avirulent *Z. tritici* strain can penetrate, colonize and reproduce asexually on a resistant host in a mixed infection with virulent strains. Microscopic observations of the disease progress indicate that virulent *Z. tritici* strains facilitate the infection and asexual reproduction of co-infecting avirulent strains. We further demonstrated following a comparative transcriptomic approach that virulent strains suppress avirulent strain-triggered host immune responses. Our findings indicate that mixed infections can promote the persistence of avirulent pathogen strains in resistant plant populations.

## Results

### Mixed infections facilitate the asexual reproduction of avirulent strains

We first investigated if avirulent strains can asexually reproduce in the presence of virulent strains. We simultaneously co-infected the Stb6-containing resistant cultivar Chinese Spring with two strains harboring either virulent (3D7 or 1A5) or avirulent (1E4) isoforms of AvrStb6. To discriminate between strains, we used eGFP-labeled 1E4 and mCherry-labeled 3D7. We observed 1E4-eGFP cirri nearly exclusively on leaves co-infected with 1A5 or 3D7-mCherry (Fig. 1A and B). Pycnidia from the avirulent strain 1E4-eGFP were rarely observed in single infections (3 pycnidia observed in 48 leaves), while the reproductive success of 1E4-eGFP was 22 times higher (67 pycnidia observed in 48 leaves; t-test, p = 0.04) in mixed infections with virulent strains (Fig. 1C, Supporting information Table S1). The pycnidia density of the virulent 3D7-mCherry strain was identical in both single and mixed infections (Fig. 1D, Supporting information Table S1). We conclude that mixed infections of avirulent and virulent strains do not affect the spore production of the virulent strain, but significantly increase the reproductive success of the avirulent strain.

**Figure 1:**
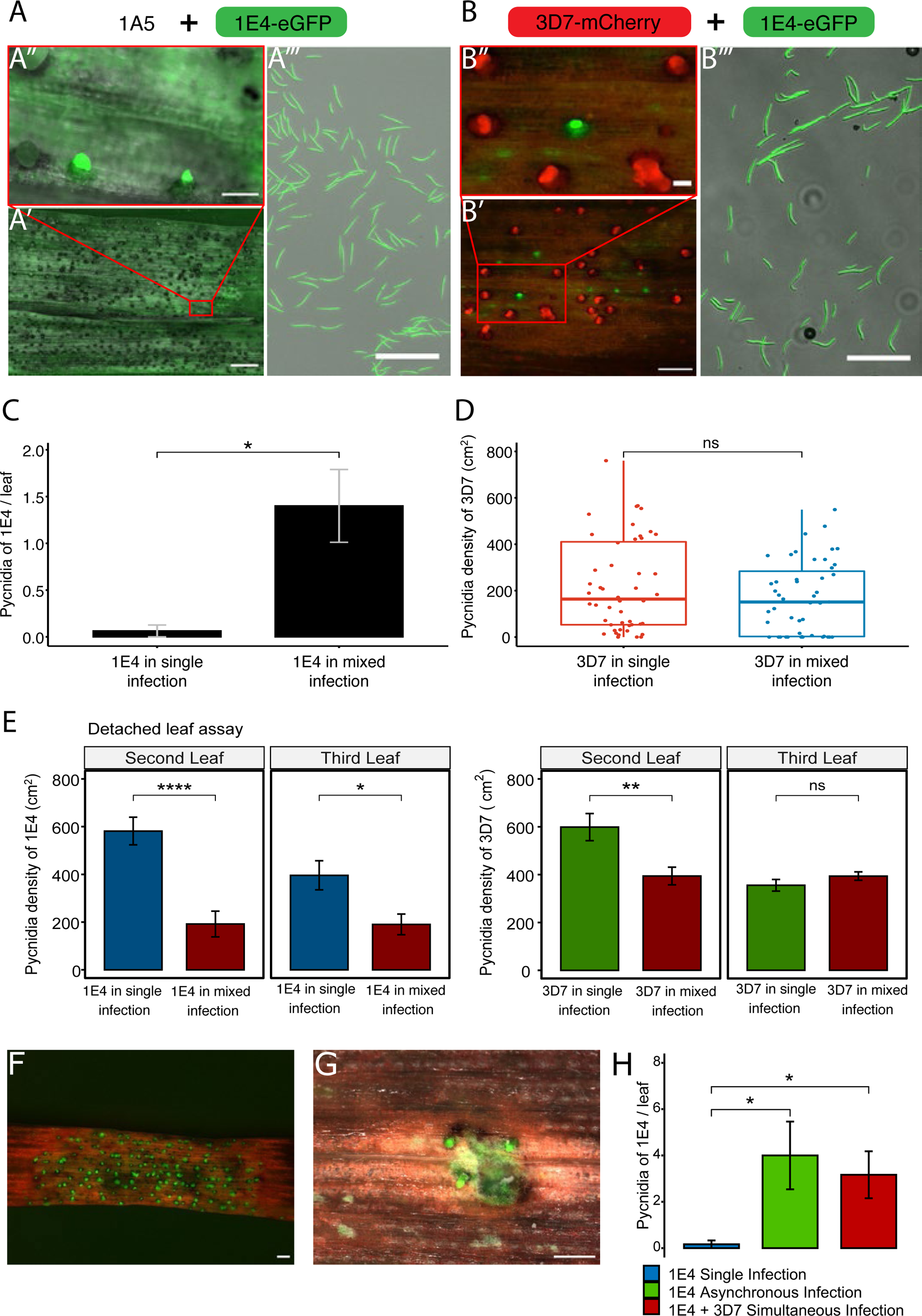
*Zymoseptoria tritici* avirulent strain 1E4 can reproduce in mixed infections with virulent strains. **A)** The avirulent strain 1E4 tagged with eGFP produces pycnidia on the resistant wheat cultivar Chinese Spring in mixed infections with the unlabeled virulent strain 1A5. Leaves were harvested at 19 dpi. A’ and A”: Leaf surface with mature fluorescent pycnidia and extruded cirri of 1E4-eGFP. Scale bar: 1 mm (A’) and 100 µm (A”). The red box in A’ indicates the zoom out area displayed in A”. A’”: Pycnidiospores collected from pycnidia presenting a fluorescent signal. Scale bar: 100 µm. **B)** The avirulent strain 1E4-eGFP produces pycnidia in the presence of the virulent strain 3D7 labeled with mCherry on the resistant cultivar Chinese spring. B’ and B”: Pycnidia distribution pattern of 3D7-mCherry and 1E4-eGFP. Scale bar: 500 µm (B’) and 100 µm (B”). The red box in B’ indicates the zoom out area displayed in B”. B’”: pycnidiospores collected from the eGFP-labeled pycnidia are from 1E4-eGFP. Scale bar: 100 µm **C)** 1E4 produces more pycnidia in mixed infections with the virulent strain 3D7 than in single infections. Asterisk indicates significant differences according to one-tailed student test (P < 0.05). **D)** The virulent strain 3D7 produces the same number of pycnidia regardless of the presence of the avirulent strain 1E4. Pycnidia of 3D7 per cm^2^ of leaf are represented. No significant differences were detected according to the Wilcoxon signed-rank test (P < 0.05). Data in B), C) and D) are from three biological replicates. A total of 48 leaves were taken for each infection experiment. **E)** Barplot of the average number of pycnidia per square centimeter leaf from 1E4-eGFP (left panel) and 3D7-mCherry (right panel) produced on detached leaves of Chinese Spring in single and mixed infections. Two cm from the leaf tip were discarded, and four pictures were taken randomly in the adjacent 17 cm of each leaf. Error bars represent the standard error of the mean. Asterisks indicate significant statistical differences according to the student test (* = p≤0.05, ** = p≤0.01, *** = p≤0.001, **** = p≤0.0001 and ns = not significant). The experiment was performed in triplicate. **F)** and **G)** Pictures of pycnidia of 1E4 labeled with eGFP at 20 days post infection (dpi) in Chinese Spring leaves damaged with celite **F)** or poked with a needle **G)** prior to inoculation. Scale bars indicate 500 µm. The experiment was performed twice. **H)** Asynchronous and simultaneous infections favor reproduction of the avirulent strain on the cultivar Chinese Spring. Barplot of the average pycnidia density per leaf of 1E4 infected alone, infected 7 days after inoculation with 3D7 (asynchronous) and infected simultaneously with 3D7. Infections were evaluated 20 days after inoculation of 1E4. Data is from three biological replicates. The experiment was performed twice. Asterisks indicate significant differences according to Anova followed by Tukey’s HSD (p < 0.05).

We hypothesized that the increase in the reproductive rate of the avirulent strain 1E4 in co-infections could be due to direct interactions between the strains, plant cell damage produced by the virulent strain or to the suppression of the immune response by the virulent strain. We first investigated how the host environment influenced the interaction between virulent and avirulent strains by performing infection assays on detached leaves (Liu *et al*., 2007). The avirulent strain 1E4-eGFP achieved significantly higher pycnidia density in single infections than in mixed infections on both second (p < 0.001) and third detached leaves (p = 0.015; Fig. 1E, Supporting information Table S2). Pycnidia density of the virulent strain 3D7-mCherry was reduced on the second leaf (p = 0.007), but not on the third leaf (p = 0.215) by the presence of 1E4-eGFP (Fig. 1E, below, Supporting information Table S2). These results indicate that asexual reproduction on detached leaves is potentially limited by competition (Barrett *et al*., 2021) between avirulent and virulent strains. In contrast, in intact leaves we observed that asexual reproduction of the avirulent strain is favored by the presence of the virulent strain (Fig. 1C). We therefore suggest that interactions between strains are conditional on the host environment, in this case facilitating asexual reproduction of avirulent strains in intact resistant plants.

We next investigated whether plant cell damage produced by the virulent strain benefited the avirulent strain by determining whether mechanical leaf damage affects the reproductive success (i.e., pycnidia production) of the avirulent strain. Chinese Spring leaves were damaged with celite or poked with a needle before infection with 1E4-eGFP. We observed that the avirulent strain produced pycnidia on the leaf regions damaged with celite and on 20% of the areas poked with a needle. However, no pycnidia of the avirulent strain were formed on the intact regions of the leaf (Fig. 1F and G). The results suggest that damage to the leaf epidermis frequently allows the fungus to penetrate the plant tissues and produce pycnidia. We further evaluated whether the necrosis induced during infection promotes the colonization by the avirulent strain by instead of simultaneously infecting both strains, infecting first Chinese Spring plants with the virulent strain 3D7-mCherry and re-infected them with the avirulent strain 1E4-eGFP seven days later. Thereby 1E4-eGFP infection was initiated during the 3D7-mCherry latent phase and only 5 days before 3D7-mCherry necrotrophic phase. As a control, we conducted simultaneous infections, in which both strains were co-inoculated as previously indicated (Fig. 1 A-C). In asynchronous co-infections, 1E4-eGFP produced more pycnidia in mixed infections than in single infections (p = 0.023; Fig. 1H). The prior infection by the virulent strain did not significantly increase the number of pycnidia of the avirulent strain, compared to the simultaneous infection (Fig. 1H, Supporting information Table S3). This suggests that the capacity of 1E4-eGFP to reproduce is affected by other factors beside the length of the latent phase and 3D7-mCherry-induced necrosis. We postulate,therefore, that other mechanisms, such as suppression of plant resistance by the virulent strain, might be involved.

### The virulent strain facilitates hyphal penetration of the avirulent strain

To test whether 1E4 is favored in mixed infections with virulent strains at the early stages of the infection, before symptoms become visible, we monitored hyphal plant penetration using confocal microscopy. Avirulent strains progression upon plant infection can be subdivided in 3 phases: stage I occurs when hyphae start to penetrate through the stomata, but do not reach the substomatal cavity; stage II occurs when hyphae reach the substomatal cavity; and stage III occurs when hyphae enter into the substomatal cavity and reach the apoplastic space in the mesophyll (Fig. 2A). At the first stages of penetration (I and II), there were no differences in the penetration rate of the avirulent strain in single and mixed infections with the virulent strain (Fig. 2A and B, Supporting information Fig. S1 and Table S4). However, the number of hyphae from the avirulent strain 1E4-eGFP that reached the apoplastic space (stage III) was significantly higher (p = 0.029) in mixed infections compared to single infections (Fig. 2B-D, Supporting information Fig. S1 and Table S4). These results indicate that host colonization by the avirulent strain at the early stages of infection is promoted by co-infections with the virulent strain. Our data suggest that co-infection enables colonization of the apoplastic space during the asymptomatic phase of the infection, leading to an increase in asexual reproduction of the avirulent strain.

**Figure 2:**
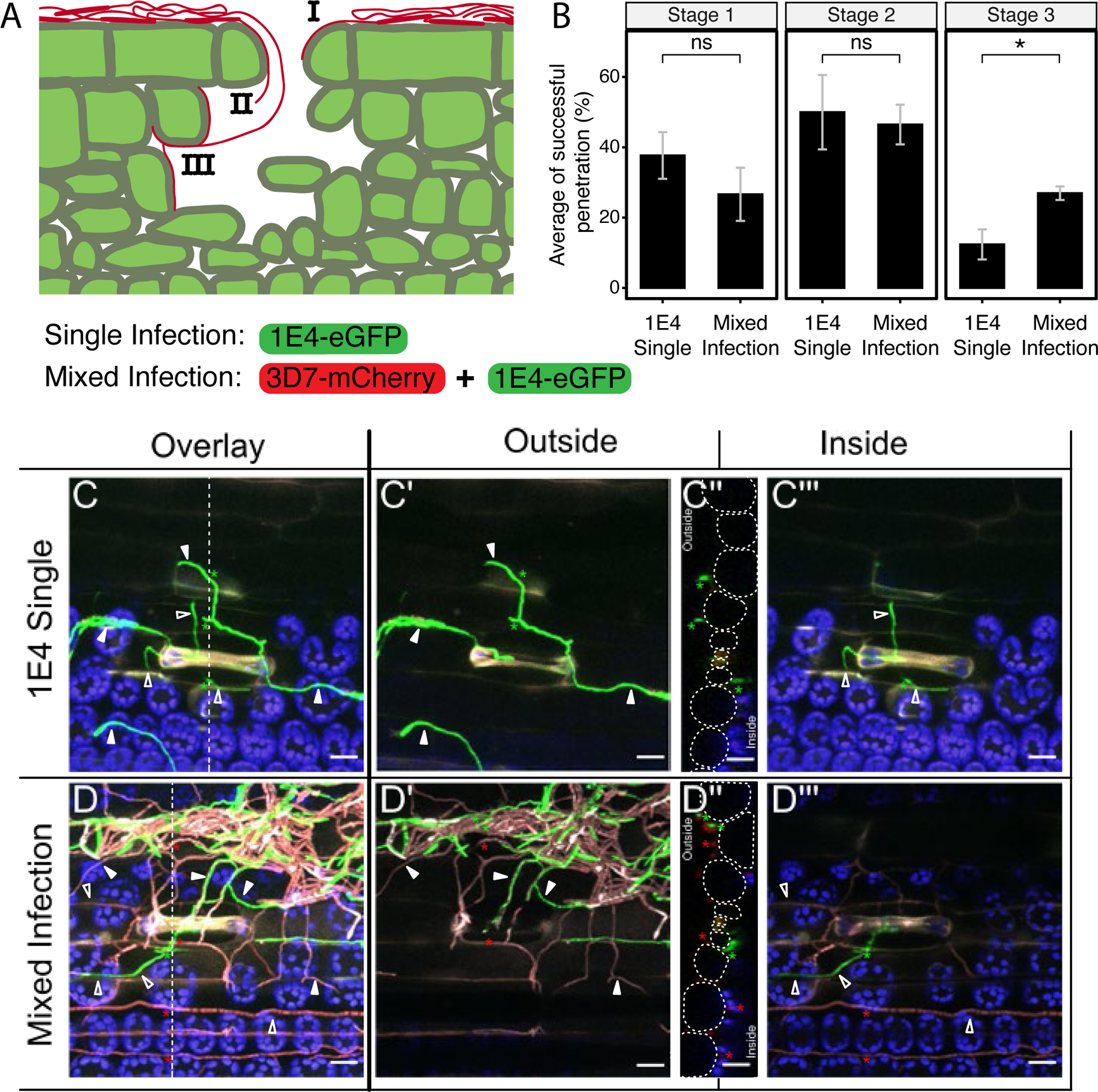
The avirulent strain reaches more frequently the mesophyll in mixed infections with a virulent strain. **A)** Penetration of the avirulent strain (1E4 labeled with eGFP) in the resistant cultivar Chinese Spring was estimated by defining four infection stages: I, the hyphae attempt to penetrate the stomata, but it does not reach the substomatal cavity; II, the hyphae reach the substomatal cavity; III, the hyphae reach the mesophyll cells. **B)** Hyphae from the avirulent strain 1E4 reach the mesophyll cells more frequently in mixed infections with the virulent strain 3D7 (labeled with mCherry) than in single infections. The penetration rate was estimated at 11 days after infection of Chinese Spring plants with 1E4-eGFP or with a mixture of 1E4-eGFP and 3D7-mCherry. Bars represent average of three biological replicates, and error bars show the standard error of the mean. In total between 10 and 29 observations per treatment and replicate were made, with a total of 64 observations made for mixed infections and 65 for 1E4 single infections. Asterisks indicate statistical differences according to two-tailed student’s test (P < 0.05). This experiment was performed twice. **C** and **D)** Maximum projection overlays of Z stack acquisitions illustrating stage III penetration events of 1E4 (1E4-eGFP, green) in single infections or in mixed infections with the virulent strain 3D7 (3D7-mCherry, red) (C and D panels respectively). Dotted lines across C and D mark the position of the orthogonal views (yz) displayed in C” and D”. Dotted circles on the orthogonal view (C”, D”) delimit epidermal cells outlines. C’ and D’ panels display the outside section from the Z stack acquisition. C’” and D’” panels display exclusively the plant inner tissues from the Z stack acquisition. Asterisks pinpoint hyphae visible on the orthogonal views. Full and empty arrowheads indicate outer and inner hyphae respectively. Chloroplast autofluorescence is displayed in blue in all C and D panels. The scale bars represent 20 µm.

### Virulent strains of *Z. tritici* manipulate the plant immune response in mixed infections

To determine if the beneficial effect of the virulent strain on the avirulent strain is systemic, we performed an infection assay in which the virulent and avirulent strains were physically separated (Fig. 3). In total, 22 pycnidia of 1E4-eGFP were counted in the leaf regions adjacent to 3D7-mCherry inoculated areas, while no 1E4-eGFP pycnidia were found on leaf sections adjacent to mock-treated areas (Fig. 3A, C). Out of the 22 pycnidia identified in regions infected with 1E4-eGFP, 10 were completely separated from the leaf region colonized by the virulent strain and 12 were located in close proximity to pycnidia of 3D7-mCherry strain (Fig. 3C). These findings indicate that infection by virulent strains of *Z. tritici* increases plant susceptibility, and this immune suppression is not restricted to the infection site, enabling infection by avirulent strains at separate locations.

**Figure 3:**
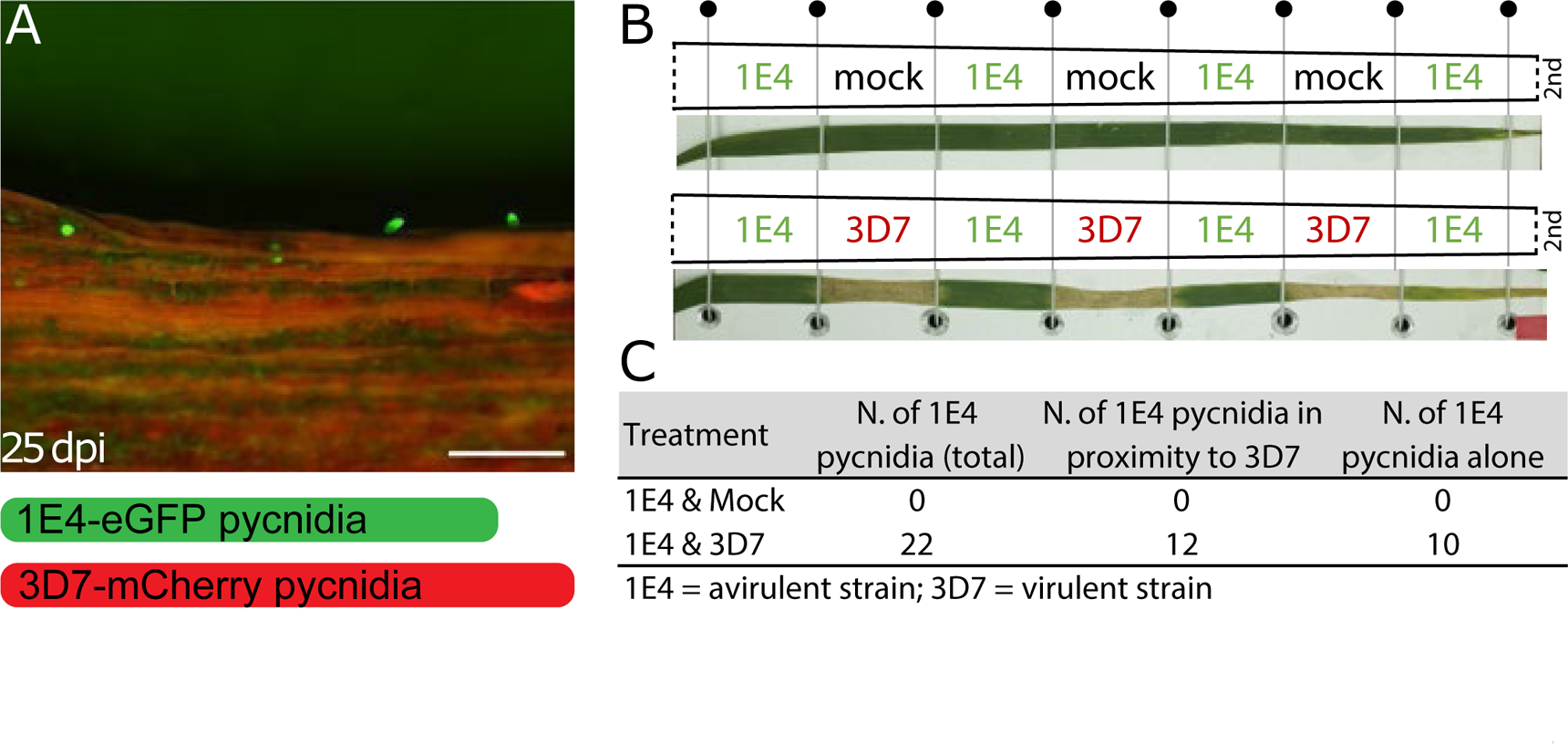
Local and distant infections favor the reproduction of the avirulent strain. **A)** The avirulent and the virulent strains were inoculated onto distant regions of the leaf. An example of pycnidia produced by 1E4 labeled with eGFP in positions distant from the virulent strain 3D7 tagged with mCherry (not in the picture). Scale bars: 500 µm **B)** Scheme of the distant infection (Side-by-side infection) performed on second leaves placed horizontally on a metal plate and kept in this position with elastic threads. Elastic threads segmented the leaf into 7 2-cm length sections. Below each scheme a representative picture of a leaves infected side-by-side with 3D7 and 1E4 or treated with the mock-solution and 1E4. Segments infected with 3D7 show necrotic lesions only between the two elastic threads, while 1E4-infected regions remain symptomless. **C)** Table summarizing the results of the side-by-side experiment indicating the number of pycnidia of 1E4 in distant infections with mock or with 3D7. The number of pycnidia from 1E4 found in the same picture as 3D7 pycnidia, or completely isolated from pycnidia of 3D7 is indicated.

To better understand the host immune response upon recognition of an avirulent strain in single and in mixed infections, we compared the transcriptomic profiles of wheat leaves infected with the virulent strain, the avirulent strain or a mixture of both strains at 3 and 6 dpi. At these time points the hyphae of both strains are mostly growing on the leaf surface and attempting to penetrate through the stomata (Haueisen *et al*., 2019). Since 1E4-eGFP alone can barely penetrate further than stage III (Fig 2B), we hypothesized that it should trigger plant immunity at earlier infection stages. We observed a clear distinction in wheat transcriptomic profiles between 3 and 6 dpi, associated with the different stages of infection (55% of variance explained on the PC1 axis; Fig. 4A). Plants infected by only the avirulent strain displayed a distinctive transcriptomic profile compared to the other infection treatments and water-sprayed leaves (mock treatment) at both 3 and 6 dpi. At these time points, the transcriptomic profile of leaves infected with the virulent strain and in mixed infections were not distinct (Fig. 4B, C), indicating that the transcriptional response of the host was independent of the presence of the avirulent strain in mixed infections. These results suggest that 3D7 overshadows the response of the avirulent strain.

**Figure 4:**
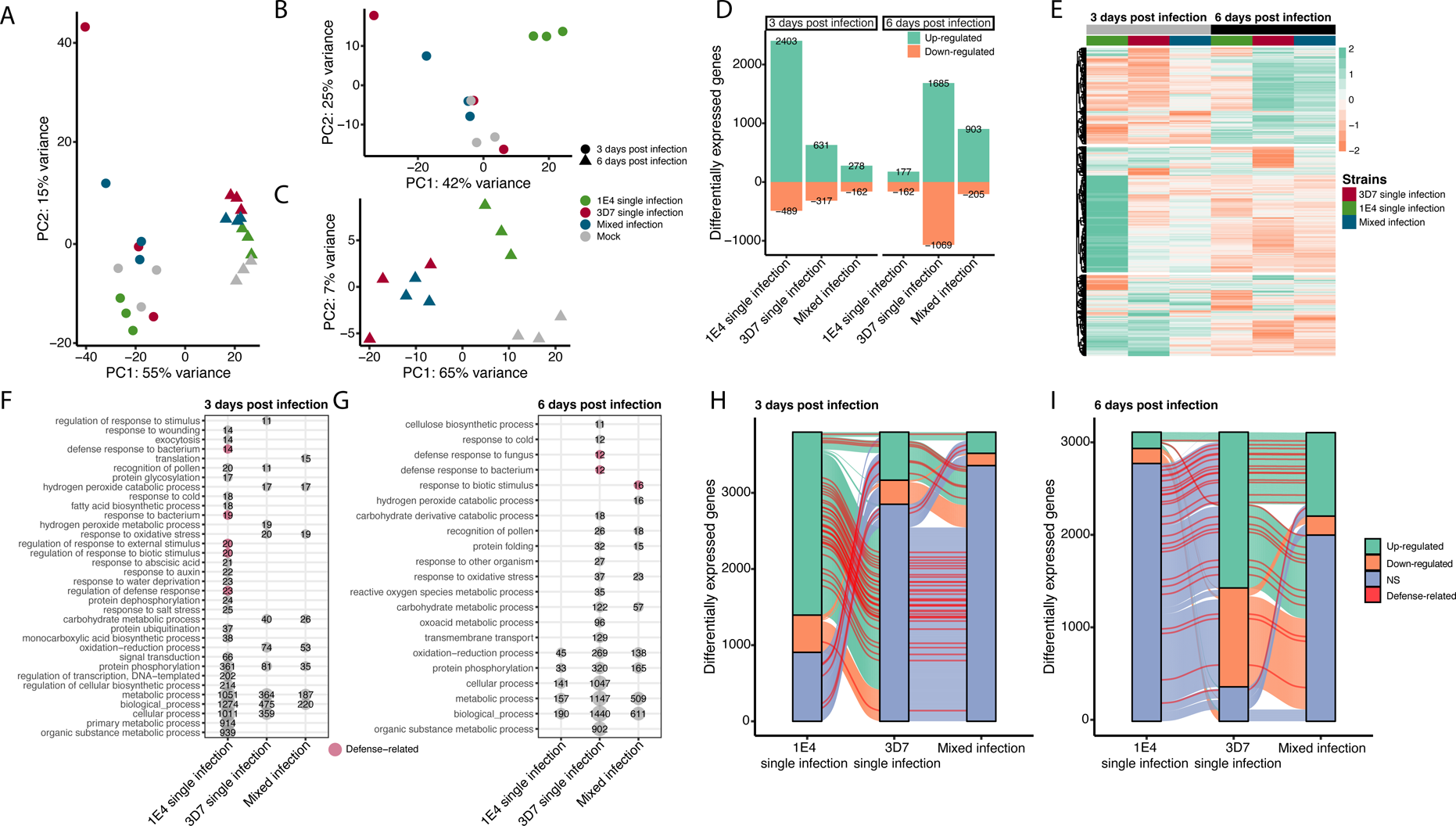
Transcriptome analysis of *Zymoseptoria tritici*­-induced early wheat response indicates that the virulent strain of *Z. tritici* controls the plant resistance. Principal component analysis (PCA) of DESeq2 normalized read counts at 3 days post inoculation (dpi; triangles) and 6 dpi (circles) for the four treatments. The four treatments include: uninfected leaves (grey), leaves infected by the avirulent strain 1E4 (green), leaves infected by the virulent strain 3D7 (red), and leaves coinfected with both 1E4 and 3D7 (blue) strains. Read counts were normalized using the size factor method and rlog transformed before performing the PCA. The PC1 (x-axis) explains 55% of the variance between treatments and time-points, while the PC2 (y-axis) explains 15% of the variance. **B)** PCA including only samples harvested at 3 dpi. Samples are placed along two PC axes explaining 42% (x-axis) and 25% (y-axis) of the variance within biological replicates. **C)** PCA including only samples harvested at 6 dpi. Samples are placed along two PC axes explaining 65% (x-axis) and 7% (y-axis) of the variance within biological replicates. **D)** Number of differentially expressed genes (DEGs) of each treatment compared to uninfected leaves identified with DESeq2 (p-adjusted < 0.05). Green bars represent up-regulated DEGs (log2 fold-change > 0) and orange bars represent down-regulated genes (log2 fold-change < 0). **E)** Transcript profiling across the three treatments and two-time points of the 6256 genes differentially expressed in at least one condition compared to uninfected plants. Transcript expression is represented in row scaled log2-fold change using uninfected leaves as a reference, from up-regulated genes (green) to down-regulated genes (orange). DEGs are grouped by hierarchical clustering based on log2-fold change values. Gene Ontology (GO) enrichment analysis of DEGs at **F)** 3 dpi and **G)** 6 dpi, only significant GO enriched categories are shown, and “defense-relateD” GOs are highlighted in red. Numbers in circles represent GO IDs annotated, only GO categories containing at least 100 annotations in the wheat genome were displayed on the plot. **H, I**) Alluvial plot of differentially expressed genes: up-regulated in green, down-regulated in orange and non-significant DEGs in blue; at **H)** 3 dpi and **I)** 6 dpi and across each treatment. Genes with “defense-relateD” GO predictions are highlighted in red.

To further explore the plant responses in single and mixed infections, we analyzed the differentially expressed genes (DEGs) in infected wheat leaves with the virulent strain, the avirulent strain or both (mixed-infection) compared to mock-treated plants at each time point. We counted 2892 and 339 DEGs (log2-fold change > 0 and p-value adjusted < 0.05) during the infection of the avirulent strain compared to the mock-infected plants at 3 and 6 dpi, respectively (Fig. 4D). In contrast, at 3 dpi the leaves infected by the virulent strain, or the mixed infection showed fewer DEGs compared to the leaves infected by the avirulent strain, with 948 and 440 DEGs, respectively. At 6 dpi, the numbers of DEGs increased in both the virulent strain infection and the mixed infection with 2754 (with 61% of genes up-regulated) and 1108 (with 81% of genes up-regulated), respectively (Fig. 4D). Taken together, these results show distinct transcriptional responses between timepoints and infection treatments.

In accordance with the principal component analysis (Fig. 4, B and C), hierarchical clustering revealed three different expression profiles for all DEGs at 3 and 6 dpi. The overall host response expression profiles during either 3D7 single infection or mixed infection were strikingly similar and distinct from the 1E4 single infection profile (Fig. 4E). The set of up-regulated genes at 3 dpi during infection by the avirulent strain 1E4 was different from the set of genes up-regulated in the virulent or mixed infections (Fig. 4E). Among the 2403 up-regulated DEGs in response to the avirulent strain at 3 dpi, only 9.5% and 4% were also up-regulated in leaves exposed to virulent and mixed strains, respectively (Fig. 4H, Table S7). Among the 1685 up-regulated DEGs in response to a 3D7 single infection at 6 dpi, 46% were also up-regulated in the mixed infection, while only 7% were up-regulated in the infection by 1E4. These results show that a similar plant response occurs when leaves are exposed to a virulent strain in a single or mixed infection, indicating that the virulent strain masks the presence of the avirulent strain in a mixed infection.

Leaves infected with the avirulent strain 1E4 included 48 up-regulated genes at 3 dpi that have Gene Ontology annotations related to “defense response” (Fig. 4F, H, Supporting information Table S8, S9, S10, S11). Only 8 and 4 genes of the same category were up-regulated in leaves infected with the virulent strain in single or mixed infections, respectively (Fig. 4F, H, Supporting information Table S8, S9, S10, S11). We also observed enrichment of GO categories representing downstream signaling in plant immune responses such as “response to abscisic aciD”, “response to jasmonic aciD” and “response to salicylic aciD” (Supporting information Table S5) in leaves infected by the avirulent strain. At 6 dpi, 34 and 22 defense response-related genes were up-regulated in plants infected with the virulent strain 3D7 or both strains, respectively, while only 2 were up-regulated in plants infected with only the avirulent strain 1E4 (Fig. 4G, I, Supporting information Tables S6, S9, Table S10, S11). Overall, the results obtained with the transcriptomic analysis indicate that the avirulent strain triggered an early resistance response in single infections, but not in mixed infections. This indicates that virulent *Z. tritici* strains are able to suppress the immune response while colonizing the host.

## Discussion

The colonization and reproductive success of avirulent strains on resistant cultivars are severely limited in single infections in which only one strain attempts to colonize the host (Kema *et al*., 1996; Jones and Dangl, 2006; Cook *et al*., 2015). In this work we demonstrated that mixed infections provide an advantage to the growth and reproduction of avirulent strains on resistant hosts. We found that this advantageous effect begins at early stages of the infection process and occurs both locally and systemically. Our findings provide new insights into the factors that could contribute to the maintenance of diversity in pathogen populations and highlight how the complex nature of pathogen-plant interactions can challenge the development of sustainable control of wheat diseases (Komar *et al*., 2008). In mixed infections, virulent strains of *Z. tritici* can suppress the plant immune system enough to allow colonization and reproduction by otherwise avirulent pathogen strains, enabling the persistence of avirulent strains in fields planted to resistant cultivars.

The transcriptomic analyses revealed that early host defense responses activated by the avirulent strain are not induced in co-infections with a virulent strain. In gene-for-gene interactions, early detection of a pathogen has been described as the key factor affecting the efficient suppression of plant pathogens (Plett and Martin, 2018). In accordance, the avirulent strain 1E4 infection progress was arrested at the early stages of infection, during penetration to the host. We additionally found that the avirulent strain induced a higher number of defense-related genes at the early stages of the infection compared to virulent strains. For example, *Stb6* was slightly but significantly up-regulated at 3 dpi in the incompatible interaction, but was not differentially expressed in infections with the virulent strain. We expected that the avirulent strain 1E4 would be detected because of the presence of the Stb6 resistance protein in Chinese Spring (Zhong *et al*., 2017; Kema *et al*., 2018; Saintenac *et al*., 2018), regardless of the presence of the virulent strain. Instead, only 4% of the genes that were up-regulated in the incompatible interaction at 3 dpi were also differentially expressed in the mixed infection. We hypothesize that virulent strains of *Z. tritici* suppress the immune response in mixed infections, either by preventing the recognition of avirulent strains or by suppressing the resistance response triggered by avirulent strains attempting to penetrate. We suggest that this suppression facilitates infection by 3D7, but also enables infection by otherwise avirulent strains of the same species. This immune suppression should occur at early stages of the infection, since we observed an enhanced penetration rate for the avirulent strain, and it is most probably systemic since it has an effect in distant regions of the leaf. This remarkable finding is in accordance with previous reports that *Z. tritici* enables the colonization of non-pathogenic species and other components of the microbiome of wheat (Kerdraon *et al*., 2019; Seybold *et al*., 2020). Our experiments add further support to the hypothesis that successful pathogens manipulate the host immune system and influence secondary infections and colonization by other microbes, including beneficial microbes, non-pathogenic and pathogenic species, and avirulent strains of pathogens (Abdullah *et al*., 2017; Seybold *et al*., 2020). We postulate that virulent *Z. tritici* strains secrete an arsenal of effectors which manipulate the host immune response, rendering the plant more susceptible to colonization by coexisting pathogens and other microorganisms. We believe that, similar to other pathogens (Weiberg *et al*., 2013), host immune suppression is a general mechanism used by *Z. tritici*.

Mathematical models suggest that avirulence alleles are likely to disappear in natural populations due to the evolutionary pressure exerted by resistance genes (Brown and Tellier, 2011; Thrall *et al*., 2016; Stephens *et al*., 2021). Models also indicate that mixed infections are likely to impact the evolution of virulence (Alizon and Van Baalen, 2008). Previous work demonstrated that avirulent and virulent strains of *Z. tritici* are capable of sexually recombining on resistant cultivars, providing a mechanism to maintain avirulent *AvrStb6* alleles in *Z. tritici* populations (Kema *et al*., 2018). Our experiments showed that mixed infections of *Z. tritici* can facilitate the asexual reproduction of avirulent strains, which provides another mechanism to maintain avirulent effector alleles at low frequencies in resistant host populations. We speculate that the same mechanisms will enable avirulence alleles to be maintained in resistant host populations for many plant pathogens.

## Experimental procedure

### Plant and fungal material

All experiments were conducted with the wheat cultivar Chinese Spring (*Triticum aestivum* L., Delley Semences et Plants SA, DSP, Delley, Switzerland), which contains the resistance gene *Stb6*. Depending on the experiment, 8 or 16 seedlings were grown in square 11 x 11 x 12 cm plastic pots (Bachmann Plantec AG, Switzerland) containing peat soil (Jiffy soil substrate GO PP7, Netherlands) for 17 days prior to infection. 10-day-old plants were fertilized with 2 litres of fertilizer solution per 15 pots (2 mL L^−1^, Wuxal Universal-Dünger, Maag-Garden, Switzerland). Growing conditions were the following: 16 h of light, 70% relative humidity, temperature of 18°C during the day and 15°C during the night, and light intensity of 12 kLux.

The ST99CH_3D7 and ST99CH_1A5 (abbreviated as 3D7 and 1A5, respectively) strains of *Z. tritici* are virulent on cultivar Chinese Spring (Linde *et al*., 2002; Croll *et al*., 2013; Barrett *et al*., 2021) while ST99CH_1E4 (1E4) harbors AvrStb6 and is avirulent on Chinese Spring (Zhong *et al*., 2017). To visualize and distinguish pycnidia and hyphae of the two mixed strains, we used a 1E4 variant tagged with a codon-optimized version of cytoplasmic enhanced green fluorescent protein (eGFP) and a 3D7 variant tagged with the cytoplasmic monomeric Cherry (mCherry) (Kilaru and Steinberg, 2015; Schuster *et al*., 2015; Meile *et al*., 2018; Barrett *et al*., 2021). Spores were incubated in 50 mL yeast sucrose broth (YSB, 10 g L^−1^ yeast extract, and 10 g L^−1^ sucrose supplemented with 50 mg mL^−1^ kanamycin) for 6 days at 18°C and 120 rpm. Spore cultures were filtered through two layers of sterile gauze, pelleted at 3273 g for 15 min, and resuspended into 15 - 25 mL of sterile water. The concentration of the spore inoculum was estimated using KOVA Glasstic counting chambers (Hycor Biomedical, Inc., California). The fitness of the strains used in all experiments was analyzed by applying 3 uL drops of a dilution series 1: 100 starting from the sprayed concentration onto yeast malt sucrose agar (YMA, 4 g L^−1^ yeast extract, 4 g L^−1^ malt extract, 4 g L^−1^ sucrose and 12 g L^−1^ agar) media. Plates were assessed after 6 days of incubation at 18°C (Supplementary fig. 2).

### Whole plant spray infection assays

In all experiments, unless noted otherwise, seventeen-day-old wheat plants were spray-inoculated until run-off with either one strain (1E4, 3D7 or 1A5) or a combination of two strains (3D7+1E4 or 3D7+1A5) using 10 mL of the spore suspension containing 0.1% (v/v) Tween-20 (Sigma Aldrich) or a mock solution (water and 0.1% (v/v) Tween-20) per pot. In the sequential infection experiments plants were initially spray-inoculated with a mock solution or a spore suspension of 3D7-mCherry, and 7 days later the same plants were infected with 1E4-eGFP. As a positive control, 24-day-old plants were spray-inoculated with a spore suspension containing a mix of 1E4-eGFP and 3D7-mCherry. The concentration of the spore suspension was 10^6^ spores/mL for each strain, except for the confocal microscopy experiments in which 10^7^ spores/mL was used. After inoculation, pots were enclosed within a plastic bag for 72 h to ensure 100% relative humidity. Twenty days after inoculations the second and third leaves were harvested and used for pycnidia quantification under the fluorescence stereomicroscope. Whole plant infection experiments shown in Fig. 1A-C were performed three times.

### Side-by-side infection and mechanical damage assays

In the side-by-side infection and mechanical-damage assays the second leaf was placed horizontally on a flat surface similar to what was previously described (Karisto *et al*., 2019). Elastic threads were used to hold the leaves on the flat surface and to segment the leaves into 7 sections of 2 cm length. In the side-by-side infection experiments each section was treated with either 1E4-eGFP or 3D7-mCherry. In control plants, mock solution was used instead of 3D7-mCherry, adjacent to 1E4-eGFP treated sections. In the mechanical damage assays leaves were pierced with a needle (Sterican Brown 0.45×12 mm BL/LB) every 1 cm along the second leaf or damaged by applying with a paintbrush a water solution containing celite (10 g/l) to a 7 cm section of the second leaf. Undamaged leaves were also included in the experiment. A spore suspension of 10^7^ spores/mL of each strain containing 0.1% (v/v) of Tween-20 or a mock solution were applied to the adaxial side of the leaves with a paintbrush, carefully avoiding contact with the elastic threads separating each leaf segment. Treated leaves were collected at 25 dpi for the side-by-side experiments and at 20 dpi for the mechanical damage experiments. Pycnidia of 1E4-eGFP were quantified by taking pictures with a fluorescence stereomicroscope. In the side-by-side experiment, we differentiated “pycnidia in proximity to 3D7” when in the same picture, pycnidia from both strains were identified. “Pycnidia of 1E4 alone” were those that did not have any visible pycnidia of 3D7 in the proximity. These experiments were performed two times.

### Detached leaves infection assay

The top 2 cm of leaves of the 24-day-old plants were discarded and the adjacent 17 cm sections were placed flat onto water agar (1%) supplemented with 50 mg mL^−1^ kanamycin to keep the leaves moist. Spore suspensions of 1E4-eGFP, 3D7-mCherry, or a 1:1 mixture (10^6^ spores/mL for each strain) were applied with a paintbrush. Agar plates with the leaves were kept at the growing conditions described earlier for 12 days. At this stage, pycnidia were visible. Four images from random areas of each leaf were obtained with the fluorescence stereomicroscope and pycnidia of each strain were quantified manually.

### Visualization of oozing pycnidia and pycnidiospores

To visualize and quantify pycnidia, plants were analyzed between 19 and 25 dpi. Since mature pycnidia are heavily melanized, GFP emission can only be observed from oozing cirri. The first 2 cm from the leaf tip were discarded, and the adjacent 5 cm section was placed on YMA amended with 50 mg mL^−1^ kanamycin. The leaves were incubated at 18°C with 100% humidity for 24 hours to stimulate pycnidia oozing. A Leica M205 FCA stereomicroscope equipped with a Leica DFC 7000 T CCD color camera was used to capture images (Software Leica Application Suite X). The following filters were used to detect signal emissions: ET GFP (525 nm to 550 nm) and ET mCherry (630 nm to 675 nm). LED light without filters was used to obtain images on the bright field. The Fiji package from ImageJ (Version 2.3.0/1.53f, https://fiji.sc) was used to extract the pictures from the lif format and transform the images into jpg format. Pycnidiospores from single 1E4-eGFP pycnidia were collected with a sterile needle and dispersed into a droplet of sterile water placed on a microscope slide. A Leica DM2500 fluorescence microscope equipped with a Leica DFC3000 G gray-scale camera (Leica Microsystems, Wetzlar, Germany) and the filter blocks for GFP (480/40 nm excitation, 527/30 nm emission) and mCherry (580/20 nm excitation, 632/60 nm emission) were used to observe the pycnidiopores. The reproduction capacities (pycnidia quantification) of the virulent and the avirulent strains in single and in mixed infection were estimated.

### Confocal laser scanning microscopy

Confocal laser scanning microscopy was performed on an inverted Zeiss LSM 780 confocal microscope using two illumination sources, DPSS (561 nm) and Argon (488 nm) lasers. Signal detection for eGFP (494.95 - 535.07 nm); mCherry (625.61 - 643.42 nm) and chloroplast autofluorescence (656.01 - 681.98 nm) were set. Eleven days after infection, second leaves were collected, 3 cm of the tip of each leaf were discarded, and the adjacent section of 1.5-2 cm was mounted in 0.02% Tween-20. The entire adaxial side of the leaf segment was visually scanned under the microscope for penetration attempts. Images were processed using the Fiji package of ImageJ (Version 2.3.0/1.53f), and it included brightness and contrast adjustments, median filters (radius of 2 pixels), generation of maximum intensity z-projection, three-dimensional (3D) reconstruction, orthogonal projections, cropping, and addition of the scale bar. 3D reconstruction and orthogonal projections enabled us to localize the hyphae on the leaf section and to determine the stage of hyphal penetration (Fig. 3A and B). In total at least 10 leaves per treatment and biological replicate were observed. In each biological replicate, independent inocula of each strain and pots were used. Before confocal observation, a photo of each leaf segment was collected to show the absence of lesions. The experiment was repeated twice.

### RNA isolation and sequencing

Wheat plants were infected with 1E4-eGFP, 3D7-mCherry, the combination of both strains or with a mock solution. For each treatment, two leaves were pooled and a total of three biological replicates were obtained at 3 and 6 dpi. Three cm from the tip were discarded, and 6 cm of the adjacent leaf were flash-frozen in liquid N_2_ and homogenized with zirconium beads (1.4 mm diameter) using the Bead-Rupture equipped with a cooling unit (Omni International, Kennesaw, GA, USA). RNA isolation was performed using GENEzol reagent (Geneaid Biotech, Taipei, Taiwan) following the manufacturer’s instructions. RNAeasy Mini Kit (Qiagen GmbH, Hilden, Germany) was used to purify the RNA, and DNA contamination was removed using the on-column DNase treatment of the RNase-Free DNase Set (Qiagen GmbH, Hilden, Germany). Ribosomal RNA was depleted by poly A enrichment, and DNA libraries were sequenced with Illumina NovaSeq 6000 using 150 bp paired-end reads. A total of 301.7 Gb of raw reads were produced. Adapters and low-quality reads were removed using Trimmomatic v0.35 (Bolger *et al*., 2014) with the following parameters: ILLUMINACLIP:adapters.fa:2:28:10 LEADING:28 SLIDINGWINDOW:4:28 AVGQUAL:28 MINLEN:50. All raw sequence data generated in this study have been deposited in the NCBI Sequence Read Archive under accession number GSE232243.

### Transcriptomics profiling

Quality filtered reads were mapped onto the *Triticum aestivum* IWGSC transcriptome reference using Kallisto v0.46.1 with default parameters for paired-end reads (Bray *et al*., 2016). Read counts were summarized using tximport v1.2.0 (Soneson *et al*., 2016). Differential gene expression and Gene Ontology (GO) enrichment analysis was performed with the R package DESeq2 (Love *et al*., 2014) and topGO (Adrian Alexa, 2021), respectively. Transcripts were considered to be differentially expressed compared to the controls (i.e., water sprayed plants) if DESeq2 p-value adjusted (padj) was < 0.05 and log2-fold change was > 0. Principal component analysis was performed with DESeq2 rlog-transformed normalized counts. GO annotations were retrieved from Ensembl using the R package BiomaRt (Durinck *et al*., 2009). We highlighted the GO annotations referring to defense responses into the category “defense-related genes” considering the following GO IDs: GO:0002215; GO:0002229; GO:0002679; GO:0006952; GO:0006968; GO:0031347; GO:0031348; GO:0031349; GO:0042742; GO:0050687; GO:0050688; GO:0050829; GO:0050832; GO:0051607; GO:0098542; GO:1900150; GO:1900367; GO:1900425; GO:1900426; GO:2000068; GO:2000071.

### Statistical analysis

Statistical analyses were conducted in R (R Core Team, 2020). The “mvnormtest” package from Shapiro-Wilk and the “car” package from Levene’s test were used to examine the normality of residuals and homogeneity of variance. In normally distributed data, a two-tailed student’s test (P < 0.05) was performed, except for the quantification of 1E4 pycnidia per leaf (fig. 1C) and the quantification of 1E4 successful penetration events in the substomatal cavity of Chinese Spring (Fig. 2B), in which a one-tailed student’s test was performed. Analysis of variance (ANOVA) followed by Tukey’s HSD post-hoc test was used to estimate differences in the reproduction of the avirulent strain 1E4 between single infection, sequential infection, and simultaneous infection (mixed infection). Differences between pycnidia quantification of the virulent strain 3D7 in single and in mixed infection on healthy plants were tested using the Wilcoxon signed-rank test. Counts of 1E4 pycnidia in the detached leaf experiment and in the sequential experiment were root square transformed to fulfill normality and homoscedasticity of residuals.

## Acknowledgments

We thank the Institute of Molecular Plant Biology at ETH Zurich for letting us access the fluorescence stereomicroscope. Confocal laser scanning microscopy experiments were performed at the Scientific Center for Optical and Electron Microscopy (ScopeM), ETH Zurich. Sample preparation for RNA extraction and the quality check were performed in collaboration with the Genetic Diversity Centre (GDC), ETH Zurich. We thank Sreedhar Kilaru and Gero Steinberg, who provided us with the fluorescent *Z*. *tritici* strains (1E4-eGFP and 1E4-mCherry). We are grateful to Cristian Carrasco for providing feedback on the manuscript. This project was supported by the ETH research grant ETH-23 15-2 to ASV and BAM and by the RYC2018-025530-I grant from the Spanish Ministry of Science, Innovation and Universities to ASV.

## Authors contribution

AB, CL and ASV contributed to writing the manuscript; JA and BAM contributed to revising the manuscript; AB, CL, JA and ASV contributed to designing the experiments, and analyzing and interpreting the data; AB, PF and JA contributed to acquiring the data; AB, CL, JA and ASV contributed to the conception of the work.

## Data availability statements

Source data for figures are provided in the supplementary tables. RNA sequencing data have been deposited in GEO NCBI, accession number GSE232243.

---

**Supplementary figure 1: The avirulent strain reaches the mesophyll in mixed infections with a virulent strain**. Hyphae from the avirulent strain 1E4 reach the mesophyll cells more frequently in mixed infections with the virulent strain 3D7 (labeled with mCherry) than in single infections. The penetration rate was estimated at 11 days after infection of Chinese Spring plants with 1E4-eGFP or with a mixture of 1E4-eGFP and 3D7-mCherry. Bars represent the average of three biological replicates, with standard error. Asterisks indicate statistical differences according to two-tailed student’s test (P < 0.05). The results are from an independent repetition of the experiment shown in Fig. 2B. The four infection stages are described in Fig. 2A.

**Supplementary figure 2: Fitness assay of the strains used in the infection assays**. Phenotypes of the strains used for the confocal microscopy infection assays in solid media. For each strain, two drops of three µL of fungal spore suspensions at a concentration of 10^6^, 10^5^ and 10^4^ spores·mL^-1^ per strain (3D7 and 1E4) were inoculated on yeast-malt-sucrose agar (YMA) and incubated at 18°C for 5 days.

**Table S1: Raw data of figure 1C-D.**

**Table S2: Raw data of figure 1E-H.**

**Table S3: Raw data of figure 2 (Table S3.1) and of supplementary figure 1 (Table S3.2).**

**Table S4: Raw data of figure 3.**

**Table S5: Biological process GO enrichment analysis of up-regulated genes at 3dpi.**

**Table S6: Biological process GO enrichment analysis of up-regulated genes at 6dpi.**

**Table S7: Differentially expressed genes in at least one condition in comparison to non-infected plants. Stb6 and Stb16q are highlighted in red and blue, respectively.**

**Table S8: Defense-related DEGs in 1E4 infection 3dpi or 6dpi, with functional annotations.**

**Table S9: Defense-related DEGs in 3D7 infection 3dpi or 6dpi, with functional annotations.**

**Table S10: Defense-related DEGs in mixed infection 3dpi or 6dpi, with functional annotations.**

**Table S11: Number of DEGs containing ‘defense-related’ annotations from Fig. 4H**

