## Supplementary figures and images for "*Zymoseptoria tritici* suppresses the host immune response and facilitates the success of avirulent strains in mixed infections"

### Supplementary Figure 1

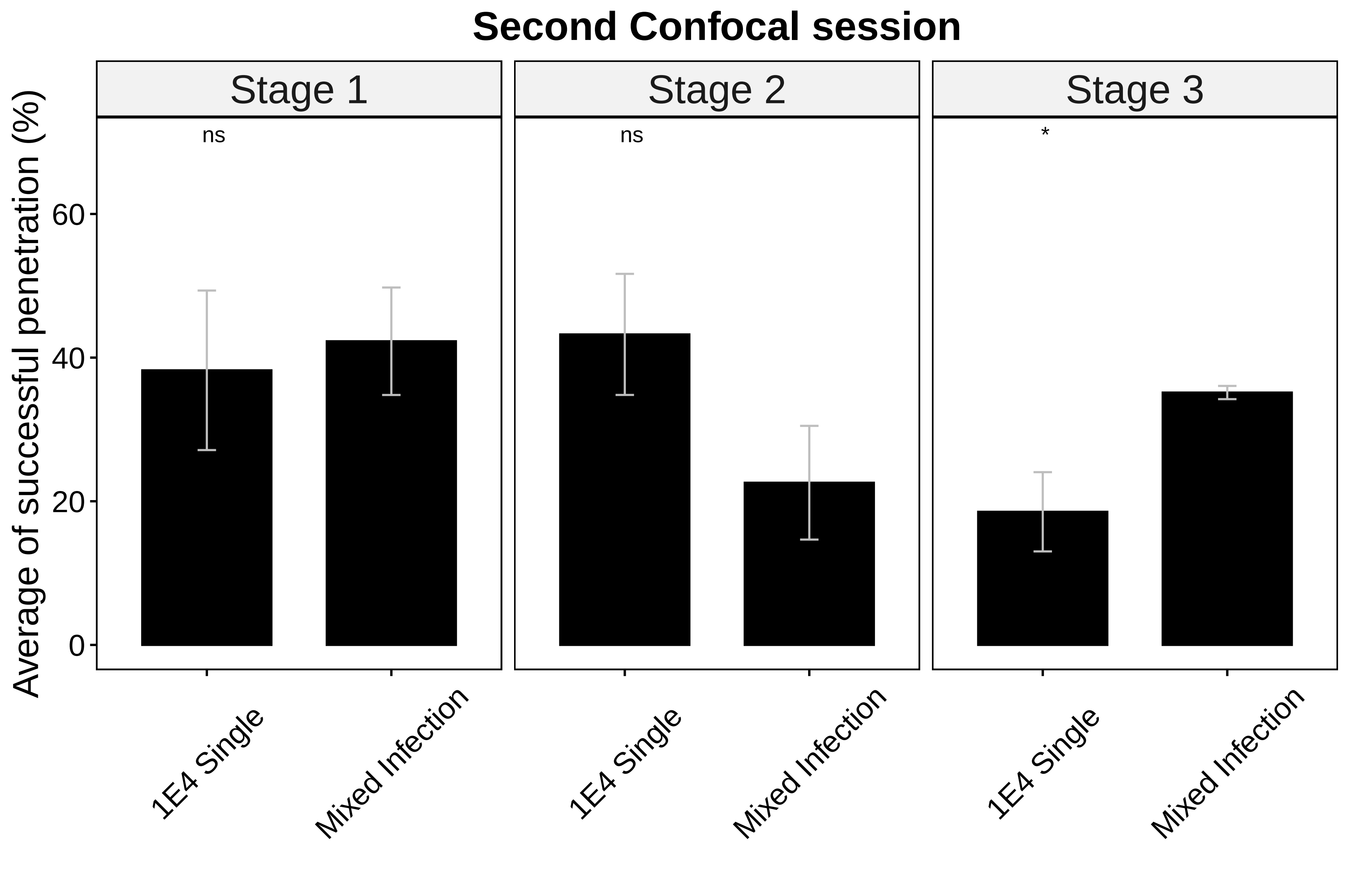

### Supplementary Figure 2

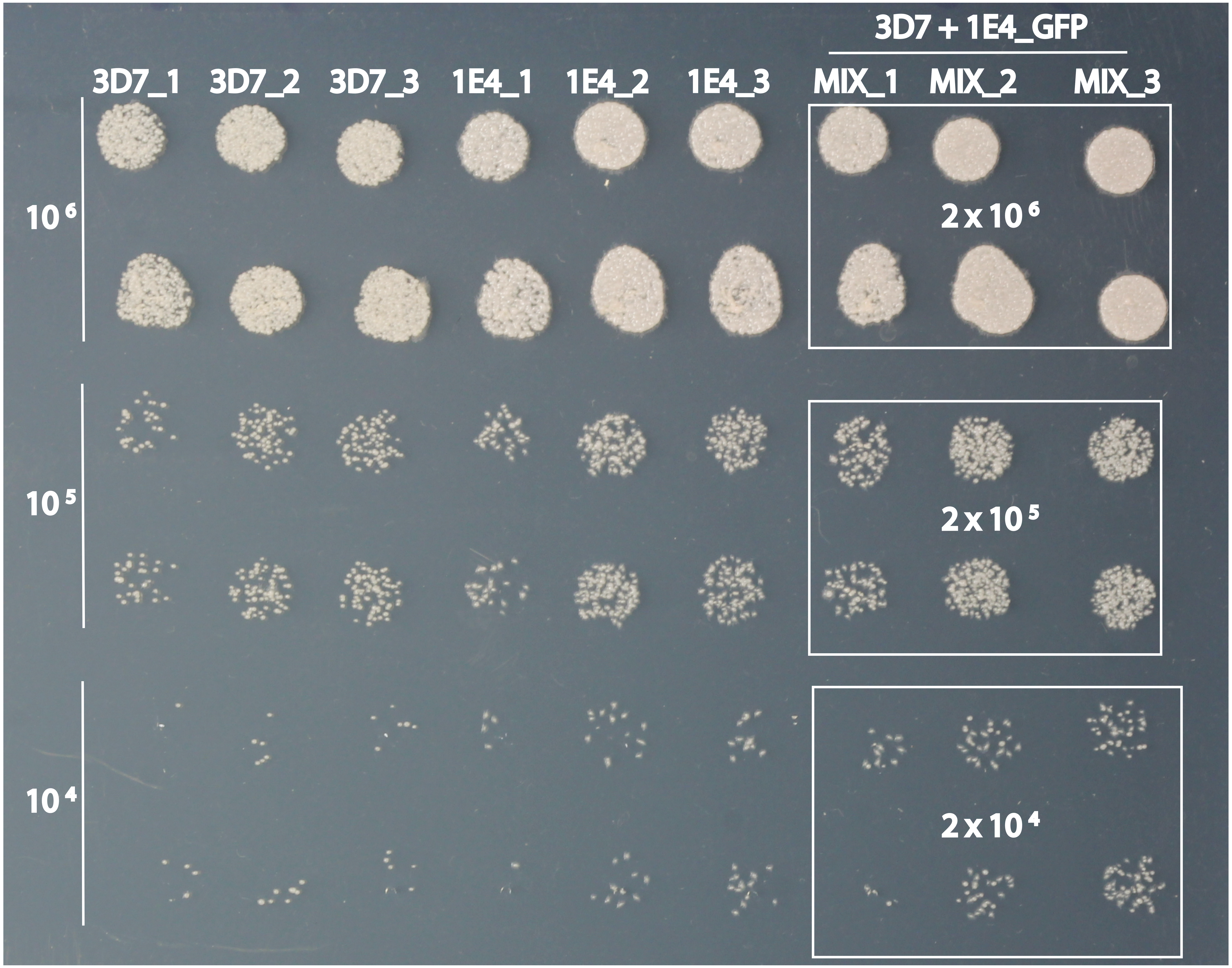
